# Open RGB Imaging Workflow for Morphological and Morphometric Analysis of Fruits using AI: A Case Study on Almonds

**DOI:** 10.1101/2025.05.05.652179

**Authors:** Mas-Gómez Jorge, Rubio Manuel, Dicenta Federico, Martínez-García Pedro José

## Abstract

High-throughput phenotyping is addressing the current bottleneck in phenotyping within breeding programs. Imaging tools are becoming the primary resource for improving the efficiency of phenotyping processes and providing large datasets for genomic selection approaches. The advent of AI brings new advantages by enhancing phenotyping methods using imaging, making them more accessible to breeding programs. In this context, we have developed an open Python workflow for analyzing morphology and heritable morphometric traits using AI, which can be applied to fruits and other plant organs. This workflow has been implemented in almond *(Prunus dulcis*), a species where efficiency is critical due to its long breeding cycle. Over 25,000 kernels, more than 20,000 nuts, and over 600 individuals have been phenotyped, making this the largest morphological study conducted in almond. As result, new heritable morphometric traits of interest have been identified. These findings pave the way for more efficient breeding strategies, ultimately facilitating the development of improved cultivars with desirable traits.

## 1. Introduction

High-throughput phenotyping is becoming a valuable tool for plant breeders and researchers to meet the challenges of future food demand (Kim, 2020; Singh et al., 2016). Breeding programs now operate in a scenario where genotype data has expanded exponentially due to the accessibility of next-generation sequencing (NGS) technologies (Bhat et al., 2016; Wetterstrand, 2013). Consequently, the implementation of genomic selection approaches and candidate gene discovery has advanced significantly (Bhat et al., 2016; Yang et al., 2020). However, these approaches also depend on phenotyping, which remains the primary bottleneck in breeding programs (Yang et al., 2020). To overcome this bottleneck, classical phenotyping tasks, which are often tedious, time-consuming, and expensive, are being replaced. Breeders are leveraging tools such as computer vision (Moore et al., 2013), artificial intelligence (Sheikh et al., 2024), and unmanned aerial platforms (Condorelli et al., 2018), among others, to meet the need for large-scale phenotyping and the generation of accurate, accessible big datasets (Yang et al., 2020).

The use of high-throughput phenotyping platforms for fruit species such as apple, mango, vineyard or citrus have helped to study architecture parameters, pigment and nutrient contents, water stress, biochemical parameters of fruits and disease detection (Huang et al., 2020). A device for fast and accurate phenotyping of small fruit samples called “FruitPhenoBox” was developed for breeding programs (Kirchgessner et al., 2024), leading to the identification of key marker-trait associations in apple populations (Keller et al., 2024). Additionally, by integrating genotype data, RGB imaging, and autoencoders, a framework was developed to reconstruct apple samples using only genotype data (Jurado-Ruiz et al., 2023). Morphometric approaches using machine learning have been applied in strawberries to extract quantitative shape features, enabling the identification of new heritable traits (Feldmann et al., 2020). Furthermore, an automatic pipeline to phenotype morphological traits was developed for strawberries using computer vision techniques (Zingaretti et al., 2021) and 3D imaging (Feldmann & Tabb 2022), with applicability to other fruits.

For stone fruit trees species, such as almond (*Prunus dulcis*) the implementation of these tools is still scarce. Almond, as other *Prunus* species such as apricot, present a long and expensive breeding cycle, in general, due to the long juvenility period and the timing to obtain a new improved cultivar, around 12-15 years. Therefore, increasing efficiency is a key aspect in breeding these species, particularly in phenotyping processes of key traits of interest as kernel morphology, shape and color (Janick & Schirra, 1997; Martínez-García et al., 2019; Socias et al., 2008). These traits directly influence market value and consumer acceptance (Demir et al., 2019). Preliminary efforts in imaging analysis and deep learning in almond have demonstrated that the same quantitative trait loci (QTLs) for size and shape can be successfully identified using these approaches as with traditional manual phenotyping methods (Pérez de Los Cobos et al., 2024). Moreover, 3D imaging phenotyping methods have been developed for almonds nuts (Sánchez-Beeckman et al., 2024).

In most of the mentioned studies, segmentation — defined as the process of partitioning a digital image into meaningful regions of interest (ROIs) based on homogeneous visual features such as color, texture, or intensity — relies on computer vision algorithms, which often require manual adjustments, limiting automation—especially as scene complexity increases (Ngugi et al., 2021; Sapkota et al., 2024). To enhance efficiency and automation, advanced tools capable of handling this complexity are essential for fruit morphology studies (Xue et al., 2024). In this sense, deep learning techniques have gained significant relevance due to their high segmentation accuracy, even in complex scenarios, by leveraging large training and validation datasets (Katal et al., 2022). Instance segmentation using deep learning can be implemented through two main approaches: two-stage models (e.g., Mask R-CNN, Faster R-CNN) and one-stage models (e.g., *You look only once* (YOLO), RetinaNet) (Gu et al., 2022). While two-stage models offer higher accuracy at the cost of lower speed and greater computational requirements, one-stage models aim to balance accuracy and speed while using fewer resources (Gu et al., 2022). Notably, recent advancements in YOLO models have significantly improved their accuracy while maintaining high-speed performance (Sapkota et al., 2024). Nevertheless, training deep learning models from scratch requires large labelled datasets and substantial computational resources (Bommasani et al., 2022). In this context, fine-tuning pre-trained generalist models allows users to develop customized models without the need for extensive datasets (Bommasani et al., 2022; Kirillov et al., 2023). The integration of deep learning tools for labelling datasets has significantly accelerated this task (Kirillov et al., 2023; Sekachev et al., 2020). Moreover, new techniques such as Slicing Aided Hyper Inference (SAHI) have enhanced accuracy by reducing errors in the segmentation of small objects, making it particularly valuable for seed imagery (Akyon et al., 2022). The slicing process also reduces memory requirements for processing large images, maintaining high resolution without the need to resize them for model training, and enabling the utilization of limited GPU (Graphics Processing Unit) open resources (e.g., Google Colab).

Here, we have developed a Python-based workflow to facilitate the development and deployment of custom deep learning models for measuring morphological and morphometric traits in fruit breeding programs. The workflow includes pre-processing methods such as color and distortion correction, image segmentation, and pixel size estimation. It has been successfully applied to phenotype multiple almond breeding populations, allowing for the estimation of broad-sense heritability values. Additionally, prediction models were developed to estimate kernel thickness based on area and weight. By implementing this workflow, the study has analyzed the largest number of individuals and data points ever recorded for the almond species to date.

## 2. Material and Methods

### 2.1. Plant material

The workflow was implemented to study breeding populations of the CEBAS-CSIC almond breeding program located in the experimental field (Santomera, Murcia SE of Spain) during 2022 and 2023 seasons. 665 unique genotypes from six F1 populations and a germplasm collection were studied (Table 1). Population parents were traditional cultivars (‘Marcona’ and ‘Desmayo Largueta’), commercial cultivars (‘Antoñeta’, ‘Penta’, ‘Tardona’, and ‘Florida’) and breeding selections (‘R1000’ and ‘S4017’). The target samples were in-shell fruits and kernels of the almond descendants used. For each genotype, around 30 in-shell fruits and 30 kernels per year were sampled, photographed, and weighed

**Table 1.**
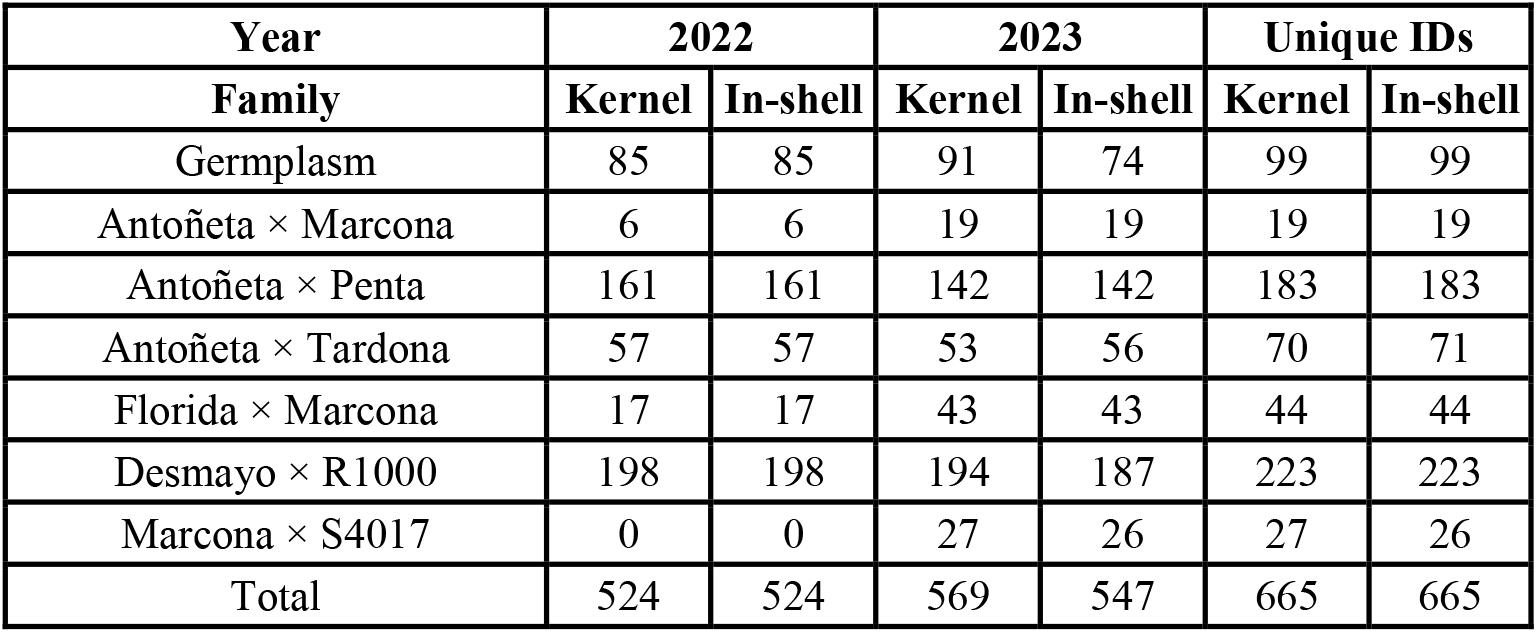
Description of the different populations used in this study, including the number of individuals per year.

| Year | 2022 |  | 2023 |  | Unique IDs |  |
| --- | --- | --- | --- | --- | --- | --- |
| Family | Kernel | In-shell | Kernel | In-shell | Kernel | In-shell |
| Germplasm | 85 | 85 | 91 | 74 | 99 | 99 |
| Antoñeta × Marcona | 6 | 6 | 19 | 19 | 19 | 19 |
| Antoñeta × Penta | 161 | 161 | 142 | 142 | 183 | 183 |
| Antoñeta × Tardona | 57 | 57 | 53 | 56 | 70 | 71 |
| Florida × Marcona | 17 | 17 | 43 | 43 | 44 | 44 |
| Desmayo × R1000 | 198 | 198 | 194 | 187 | 223 | 223 |
| Marcona × S4017 | 0 | 0 | 27 | 26 | 27 | 26 |
| Total | 524 | 524 | 569 | 547 | 665 | 665 |

### 2.2. Workflow description

A comprehensive workflow has been developed for pre-processing, segmentation model development, deployment, morphological measurements, and morphometric analyses, implemented through interactive Python notebooks (Supplementary Figure 1) (https://github.com/jorgemasgomez/almondcv2). To optimize speed and performance, the use of a GPU is highly recommended. The workflow can be executed locally; however, it is also deployed on the Google Colab platform, enabling users to run it online and leverage free resources, such as GPUs. Although primarily written in Python, some functions integrate R for morphometric analyses, requiring R to be properly configured in the system path if executed locally.

#### 2.1.1. Colour and distortion calibration

Colour and distortion correction approaches are included in the pre-processing notebook (*1_Pre-processing_workflow*.*ipynb*). Colour correction allows to standardize the dataset of the images using a colour card as reference. Colour correction functions implemented in the pre-processing notebook are based on the colour correction module of PlantCV python library (Gehan et al., 2017). Distortion correction is implemented for possible radial and tangential distortion caused by the camera. Distortion correction functions are based on the OpenCV workflow for camera calibration (*OpenCV: Camera Calibration*, 2025). It requires test patterns of chessboards with a known size of the squares to obtain a camera matrix that can be used subsequently with the pictures of the dataset. In addition, colour and distortion correction functions are joined in the pre-processing notebook to deploy it in the whole dataset in one step.

#### 2.1.2. Auxiliary functions

Some optional auxiliary functions were included for the picture pre-processing, helping to separate the different samples in the pictures and getting the physical size of a pixel with reference objects in the picture (e.g. a coin with known diameter). These optional functions require training a model (next sections) to identify and segment the sample groups and the reference objects.

#### 2.1.3. Develop your segmentation model

The approach implemented in the present work intends to simplify the development of customized segmentation models. Four key steps are necessary for performing the development: slicing, labelling, ‘training’ and reconstruction (*2_Develop_your_segmentation_model_workflow*.*ipynb*). The slicing process consists of cropping pictures into smaller patches, with the size determined by the user. Using smaller patches of high-resolution images speeds up the training and prediction processes by reducing computational requirements (Akyon et al., 2022). Additionally, this approach improves the capture of fine details, as it avoids the loss of resolution and distortion caused by resizing the entire image (Pereira et al., 2023; Saradopoulos et al., 2025). The slicing function defined in the workflow is divided into train, validation and test datasets. The slices are obtained proportionally according to the user command to be prepared for the next step. The second step, labelling, involves annotating which pixels correspond to the object of interest. For this task, the workflow has been designed to accept as input the ZIP file generated by the image annotation platform CVAT in the YOLO Segmentation 1.0 format (Sekachev et al., 2020). CVAT provides tools for semi-automatic segmentation using the Segment Anything Model (SAM) (Kirillov et al., 2023), which enables quick instance segmentation with a single click. CVAT is available for free offline via a Docker container or online with a freemium model. The third step for developing the segmentation model is ‘training’ the model. For that purpose, in the second notebook (*2_Develop_your_segmentation_model_workflow*.*ipynb*) YOLO (Redmon et al., 2016) pre-trained algorithm series can be fine-tuned (although in their web mention ‘training’ really is a fine-tuning (*Train - Ultralytics YOLO Docs*, 2025)) using our custom labeled dataset. The training process function enables to control all the parameters of the training process (e.g. epochs, batch) and provides the results of the training *YOLO*.*train* method. The last step for developing the segmentation model is to reconstruct the binary picture mask, for which two approaches were used. The first approach simply joins the patches binary masks after the prediction (*slice_predict_reconstruct*). The contours are detected using *cv2*.*findcontours* OpenCV function (Bradski & Kaehler, 2000) in the subsequent functions for morphology measurements. Watershed algorithm is implemented in such functions for separating touching objects of interest (*OpenCV: Image Segmentation with Watershed Algorithm*, 2025). The second approach uses the Slicing Aided Hyper Inference (SAHI) pipeline (*predict_model_sahi*) (Akyon et al., 2022), which is integrated in the YOLO segmentation process and the instances segmented are identified specifically providing directly their contours. Here, SAHI python library is modified slightly to enable the *retina_mask* argument in YOLO prediction function and provide fine details in the segmentations. To assess the two reconstruction approaches, there were tested in the datasets studied and the almonds with errors in the reconstruction were annotated and removed manually.

#### 2.1.4 Deploy your segmentation model for morphology and morphometric analyses

Once the segmentation model has been successfully ‘trained’ and tested, it can be deployed for morphology and morphometric analyses. In the notebook for deployment (*3_Deploy_your_segmentation_model_worflow*.*ipynb*) two methods have been prepared to measure. The first method is general for any group of fruit and the traits length, width, area, perimeter, hull-area, solidity, aspect ratio, circularity, ellipse ratio and color (L*a*b model) are measured for each fruit/seed. This general method can be used as a template and customized according to the user’s needs, for example to study additional traits or different fruit shapes. The second method is specific for almonds adding more traits such as width at three different heights (25, 50 and 75% of the length), vertical and horizontal symmetry and symmetry in the top part of the almond (to measure the shoulder). In this specific method, almonds are aligned according to the angle of fitting an ellipse (*cv2*.*fitEllipse* function) and flipped vertically with placing the most distant point always in the lowest part (almond tip). Some extra traits derived of the results together with the weight of the kernel/in-shell have been also included such as the weight kernel/in-shell ratio, estimation of the thickness (only for kernels) and globosity (width/thickness). For the estimation of kernel thickness a linear and quadratic model was fitted using 228 almond kernels, using the area and the weight as predictor variables. To validate the models the dataset was split into 80% for training 20% for validating and was run 100 times to assess the stability of the models.

Both methods export picture results, table results, and binary masks for morphometric analyses. Binary masks are flipped horizontally if the lowest pixel (tip) is the right part of the picture for avoiding symmetric issues in the morphometric analysis.

Morphometric analyses notebook (*4_Morphometrics_workflow*.*ipynb*) includes two morphometric approaches: Elliptical fourier analysis (EFA) and pixel-based principal component analysis (PCA). EFA is conducted using Momocs v1.4.1. R package (Bonhomme et al., 2014) but it has been implemented in the notebook in Python programming language executing a subprocess. Several functions are included for EFA: exploratory analysis, run the EFA, after that perform a PCA on the EFA results and *kmeans* clustering. PCA performed on EFA coefficients generate as traits the components (EF-PCs) that explain the variability of the shape. Regarding the second pixel pixel-based PCA, binary masks are flattened for performing the PCA and generate also PCs as traits (PB-PCs). Result plots of both approaches can be obtained to show the influence of each trait.

### 2.3. RGB Imaging system

For image acquisition, group pictures of various samples were captured over a black surface template, with the samples positioned inside painted rectangles. For the 2022 dataset, images were taken using a Canon EOS 70D camera, achieving a resolution of 6 px/mm. In 2023, the imaging system was redefined due to objectives beyond the scope of this work, utilizing an Arducam 8MP IMX219 camera, which resulted in images with a resolution of 4 px/mm. Both systems were illuminated using two white LED light sources.

### 2.4 Heritability

Broad sense heritability (H^2^) was estimated using the Python library *statsmodels* (Seabold & Perktold, 2010), fitting a linear mixed model using the genotype as random effects and the year as fixed effects (Equations 1 and 2).

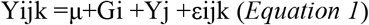

Where:

Yijk: Observed phenotypic value for the k-th observation of the i-th genotype in the j-th year.

μ: Overall population mean.

Gi: Random effect of the i-th genotype

Yj: Fixed effect of the j-th year

εijk: Residual error term

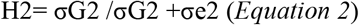

Where:

σG2 Genetic variance, which quantifies the contribution of genetic differences among genotypes to the phenotypic variance.

σe2: Residual variance, representing the combined effect of unexplained environmental variation and measurement error.

### 2.5. Impact on almond breeding

To study the impact of this new tool on almond breeding, a comprehensive comparison was conducted with existing scientific articles related to quantitative almond morphology phenotyping. Different parameters such as the number of individuals studied, sample sizes, and the number of traits analyzed were compared to evaluate the benefits of the new high-throughput phenotyping tool.

## 3. Results

### 3.1. YOLO fine-tuned segmentation models performance and reconstruction errors

A total of 46,737 elements (in-shell fruits and kernels) were segmented and performance errors from both reconstruction approaches were annotated (Table 2). A standard image generated by the workflow is shown in Figure 1. Four segmentation models were fine-tuned using datasets from kernel 2022 and 2023, and in-shell 2022 and 2023. The pre-trained model ‘yolov11s-seg.pt’ was selected, and fine-tuning was performed over 100 epochs (i.e., complete passes through the entire training dataset), using 500–1000 image slices of 320 pixels each, split into 60% for training, 20% for validation, and 20% for testing. Metrics plots from the fine-tuning process of the datasets are shown in Supplementary Figure 2 as an example Validation masks were explored, showing precise results.

**Figure 1.**
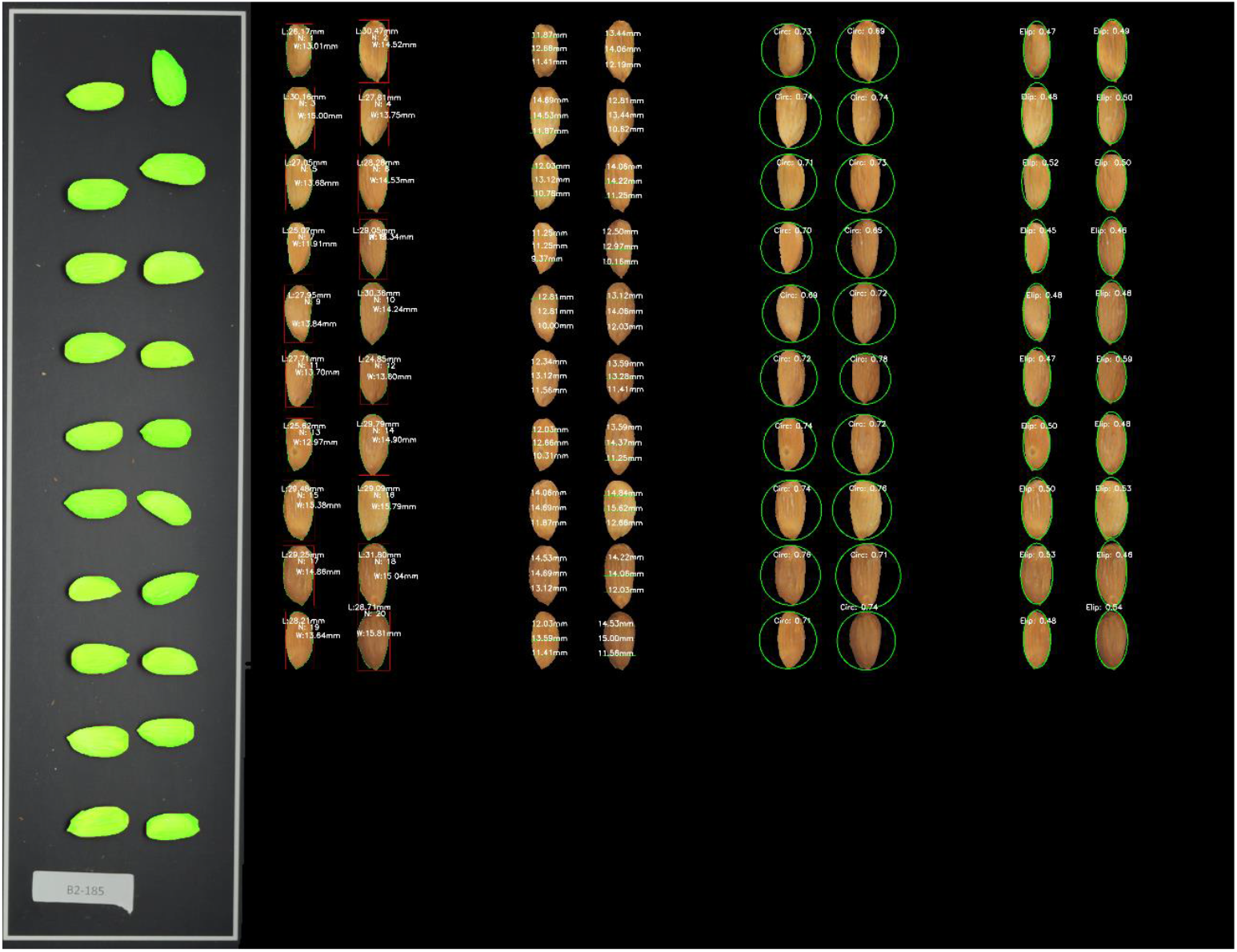
Images generated by the workflow, displayed from left to right: original image with masks, length and width measurements, widths at different lengths, circularity, and ellipse ratio.

**Table 2.** Performance summary for the different datasets and reconstruction methods, including reconstruction errors.

| Method | Dataset | Errors | Total elements | Error percentage |
| --- | --- | --- | --- | --- |
| *Slice_predict_reconstruct*<br>+<br>Watershed | Kernel-2022 | 6 | 10,180 | 0.06% |
|  | Kernel-2023 | 135 | 15,312 | 0.88% |
|  | In-shell-2022 | 4 | 10,360 | 0.04% |
|  | In-shell-2023 | 47 | 10,885 | 0.43% |
| SAHI | Kernel-2022 | 13 | 10,180 | 0.13% |
|  | Kernel-2023 | 124 | 15,312 | 0.81% |
|  | In-shell-2022 | 269 | 10,360 | 2.60% |
|  | In-shell-2023 | 43 | 10,885 | 0.40% |

All the combinations showed errors lower than 1% except in the in-shell 2022 dataset using SAHI (2.60%). Performance was similar between both approaches except a 2% difference in the in-shell 2022 dataset.

### 3.2. Thickness estimation

Kernel thickness was modelled using weight and area as predictor variables, employing both linear and quadratic models. The median R^2^ in the test dataset for the linear model was 0.725, while for the quadratic model it was 0.795. The corresponding median Root Mean Squared Error (RMSE) values were 0.56 for the linear model and 0.47 for the quadratic model (Figure 2A). The best-performing linear and quadratic models were then plotted, achieving R^2^ values of 0.847 and 0.8815, respectively, and RMSE values of 0.451 and 0.381, respectively (Figure 2B). The quadratic model trained with the complete dataset was employed for estimating thickness across all samples in subsequent analysis.

**Figure 2.**
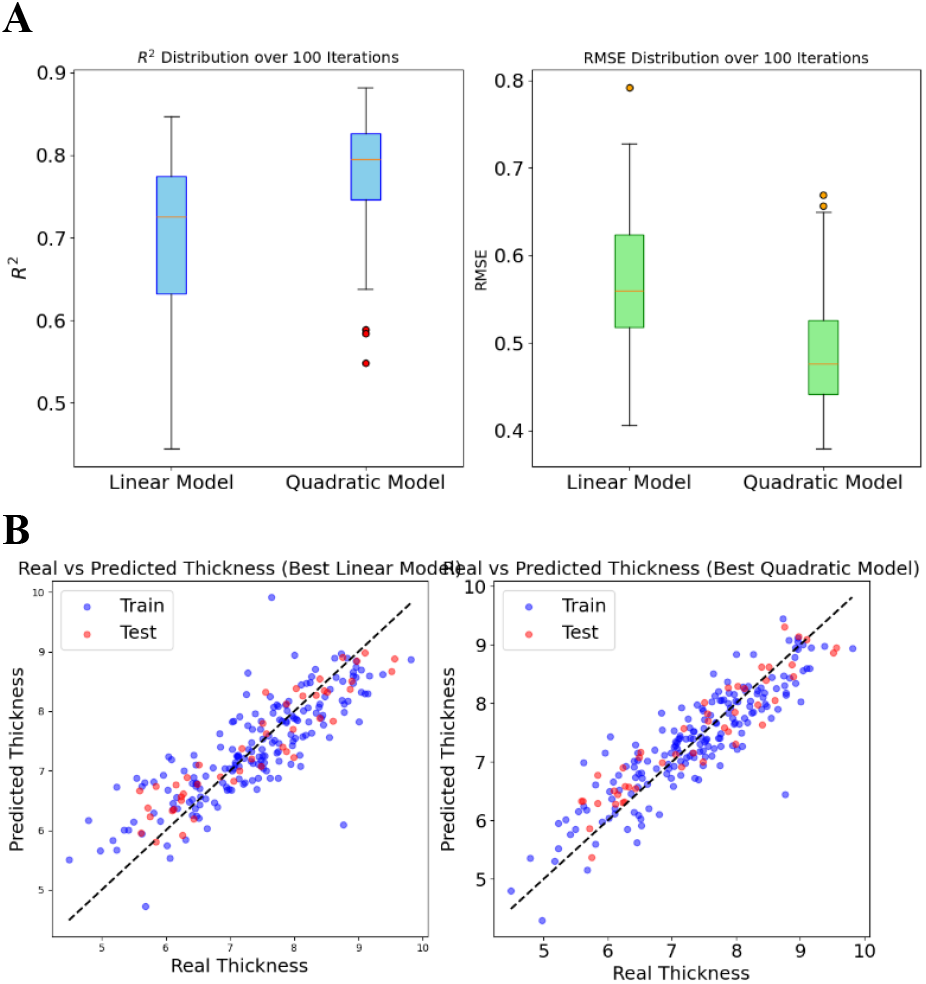
Results of the kernel thickness modeling, showing the R^2^ (A, left) and RMSE (A, right) over 100 iterations, along with scatter plots for the best linear (B, left) and quadratic (B, right) models.

### 3.3. Morphometric results

Elliptical Fourier analysis (EFA) was conducted using all the kernel and in-shell binary masks (separately) placing 10 harmonics in both datasets after a visual exploration analysis. PCA was performed on the EFA coefficients, being 62.63%, 19.93%, and 3.83% explained by the three first PCs for the kernel analysis, and 69.19%, 15.75%, and 4.15% explained by the three first PCs for the in-shell analysis (Figure 3 and Supplementary Figure 3). Only the first 6 components on each dataset were used for subsequent analysis because they explain at least 1% of the variability. K-means clustering from k=1 to k=10 was run to show the main groups of the datasets (Figure 4 and Supplementary Figure 4. The optimal number of clusters was studied and no abrupt changes in the slope by elbow method were observed in any dataset (Supplementary Figure 5).

**Figure 3.**
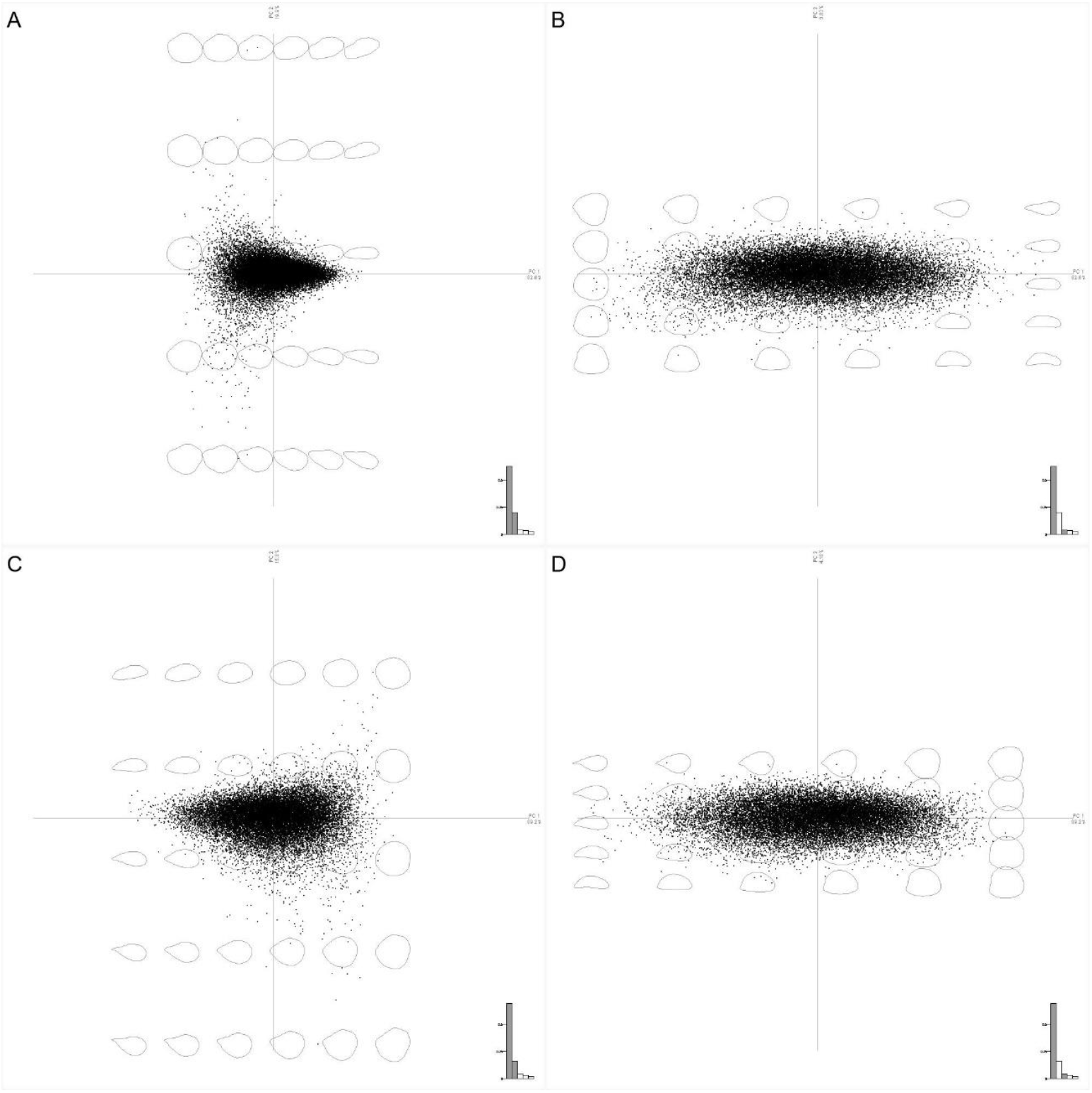
EFA-PCA scatterplot results for the kernel A) EF-PC1/EF-PC2 and B) EF-PC1/EF-PC3 and in-shell C) EF-PC1/EF-PC2 and D) EF-PC1/EF-PC3 datasets. Influence of the PCs in the shape are represented in the plots.

**Figure 4.**
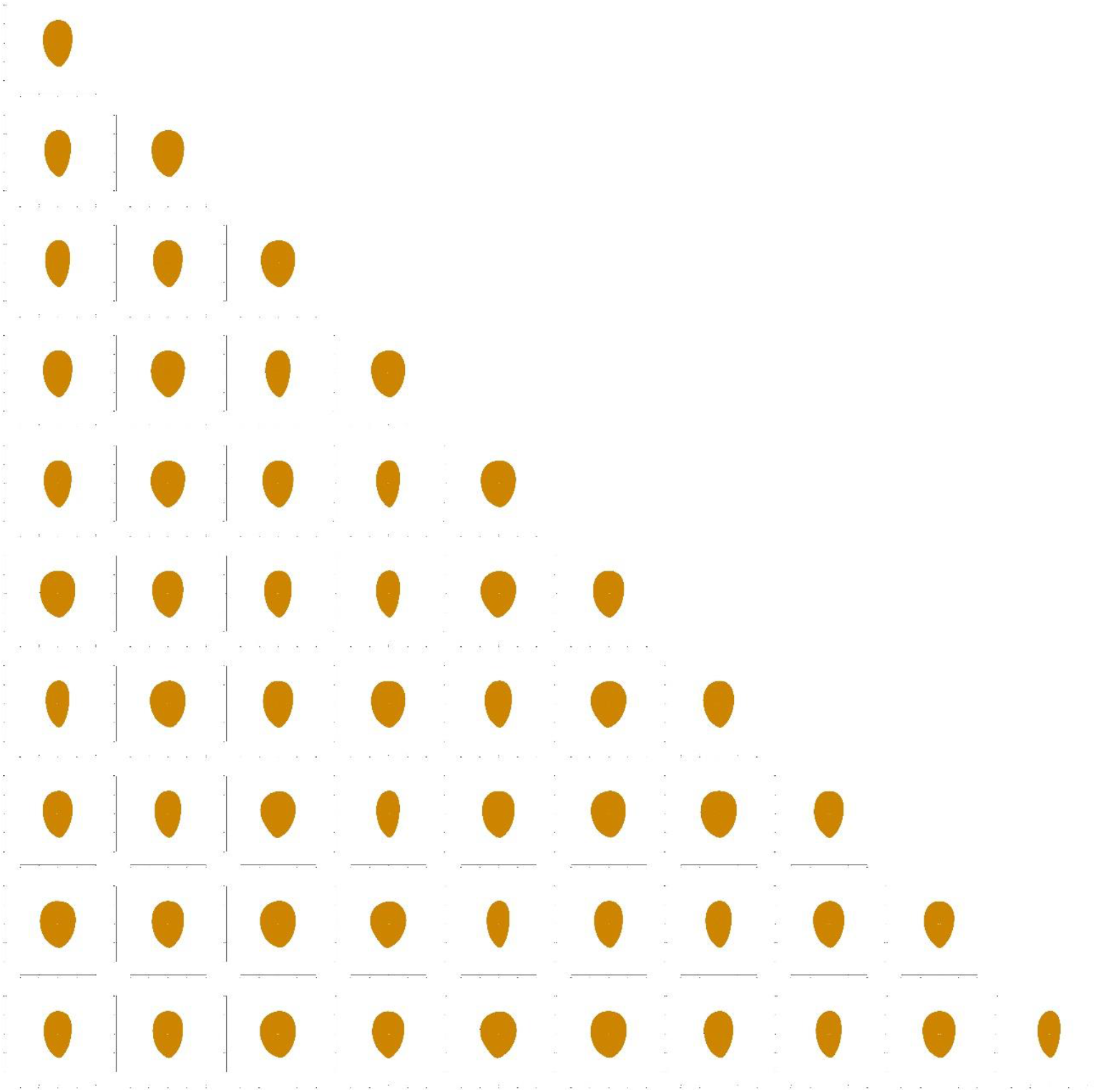
K-means clustering results using EFA-PCA in kernel dataset, showing the shapes corresponding to each centroid for scenarios ranging from k=1 to k = 10.

Pixel-based PCA was performed in kernel and in-shell datasets, obtaining 34.96%, 8.78% and 4.75% of variance explained by the three first PCs in the kernel dataset and 37.72%, 9.19% and 4.96% in the in-shell dataset (Supplementary Figure 6). The PCs influence in the shape was studied plotting the +-3x deviation from the mean shape (Supplementary Figure 7). Ten PB-PCs were collected for each dataset explaining at least 1% of variance each one of them. K-means clustering was also carried out from k=1 to k=10 (Figure 5 and Supplementary Figure 8) and no abrupt changes were observed by elbow method to determine the optimal number of clusters (Supplementary Figure 9).

**Figure 5.**
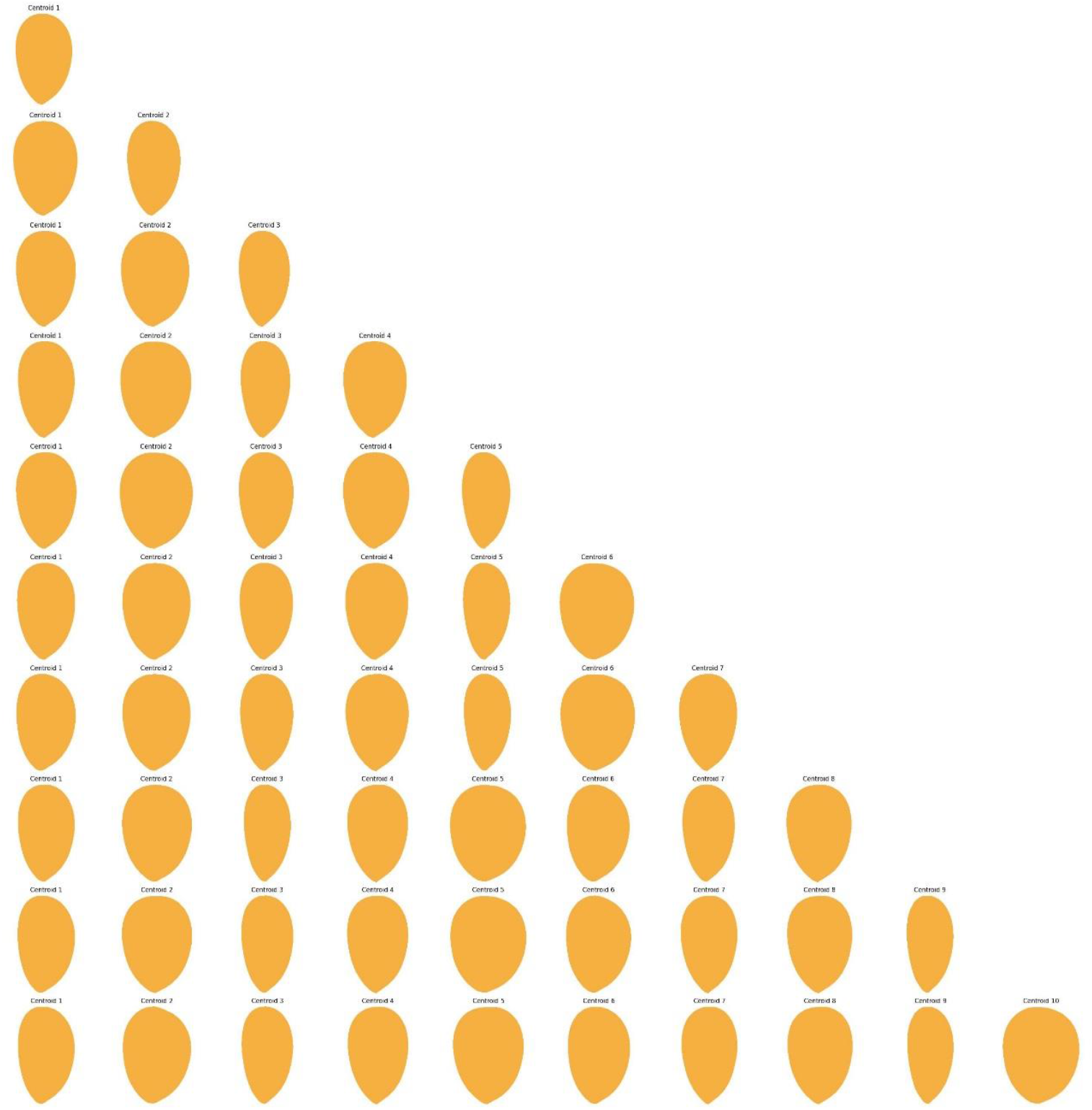
K-means clustering results using PB-PCA in kernel dataset, showing the shapes corresponding to each centroid for scenarios ranging from k=1 to k = 10.

### 3.3. Correlations

To analyse and interpret the phenotype data obtained, the Pearson correlation between all the morphological and morphometric traits was calculated (Figure 6). In absolute value, 340 pairwise correlations were higher than 0.75, with the traits of in-shell almond area, hull area, and perimeter exhibiting the highest number of strong pairwise correlations. High correlations (greater than 0.95) were observed between kernel and in-shell morphometric traits across different approaches, such as EF-PC1 with PB-PC1. Additionally, strong correlations were found between these two PC1 components and morphological traits like aspect ratio and ellipse ratio.

**Figure 6.**
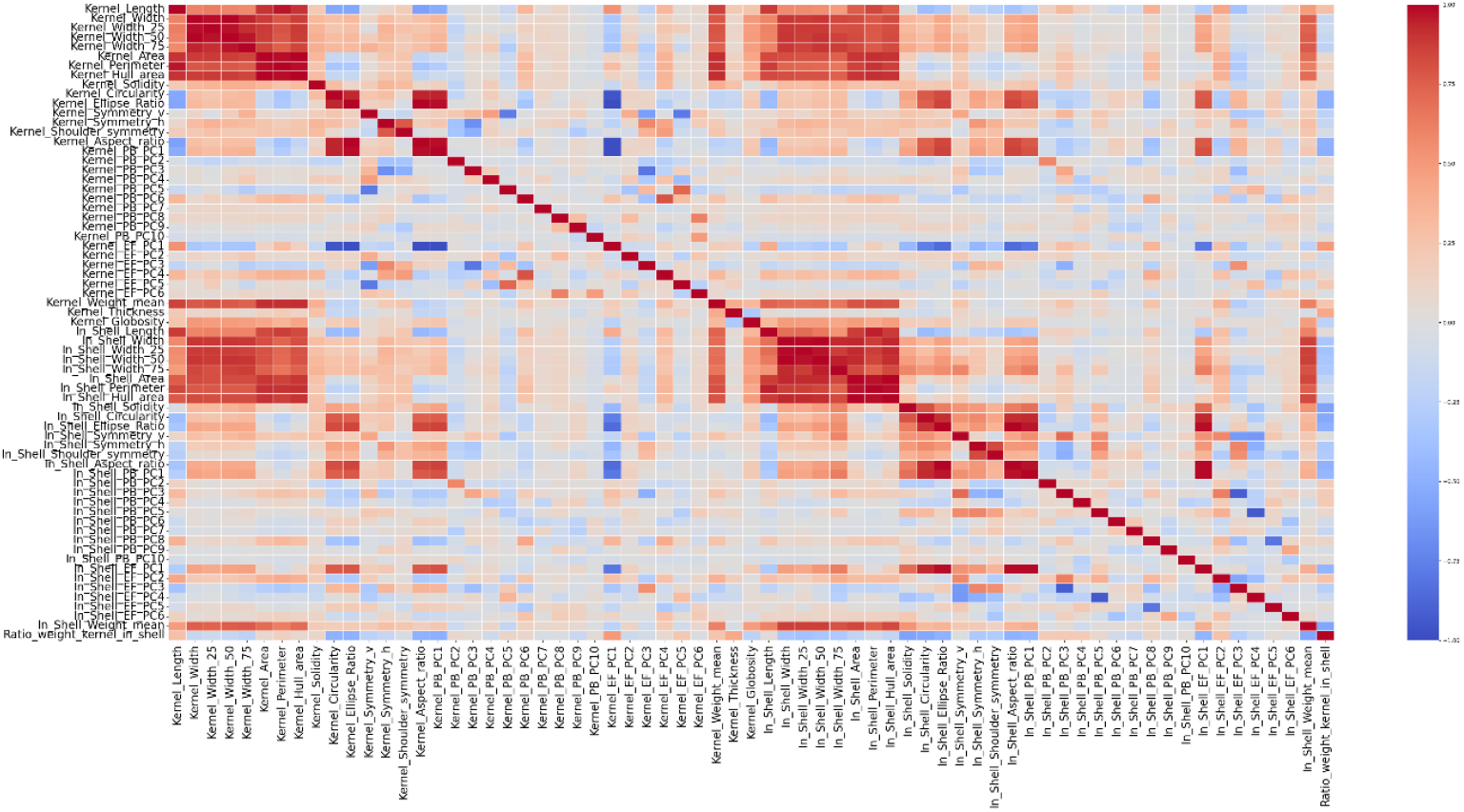
Heatmap of the Pearson correlations between all the morphological and morphometric traits.

### 3.4. Heritability

Broad sense heritability (H^2^) was estimated for all the traits studied here (Figure 7). Weight ratio kernel/in-shell, in-shell weight and in-shell EF-PC1 showed the highest heritability values (0.9, 0.77 and 0.70 respectively). Between morphometric traits, those related with PC1 showed high heritability values (higher than 0.68), and other such as Kernel PB-PC2, In-shell PB-PC2, In-shell EF-PC4 and In-shell PB-PC3 showed medium values (higher than 0.4).

**Figure 7.**
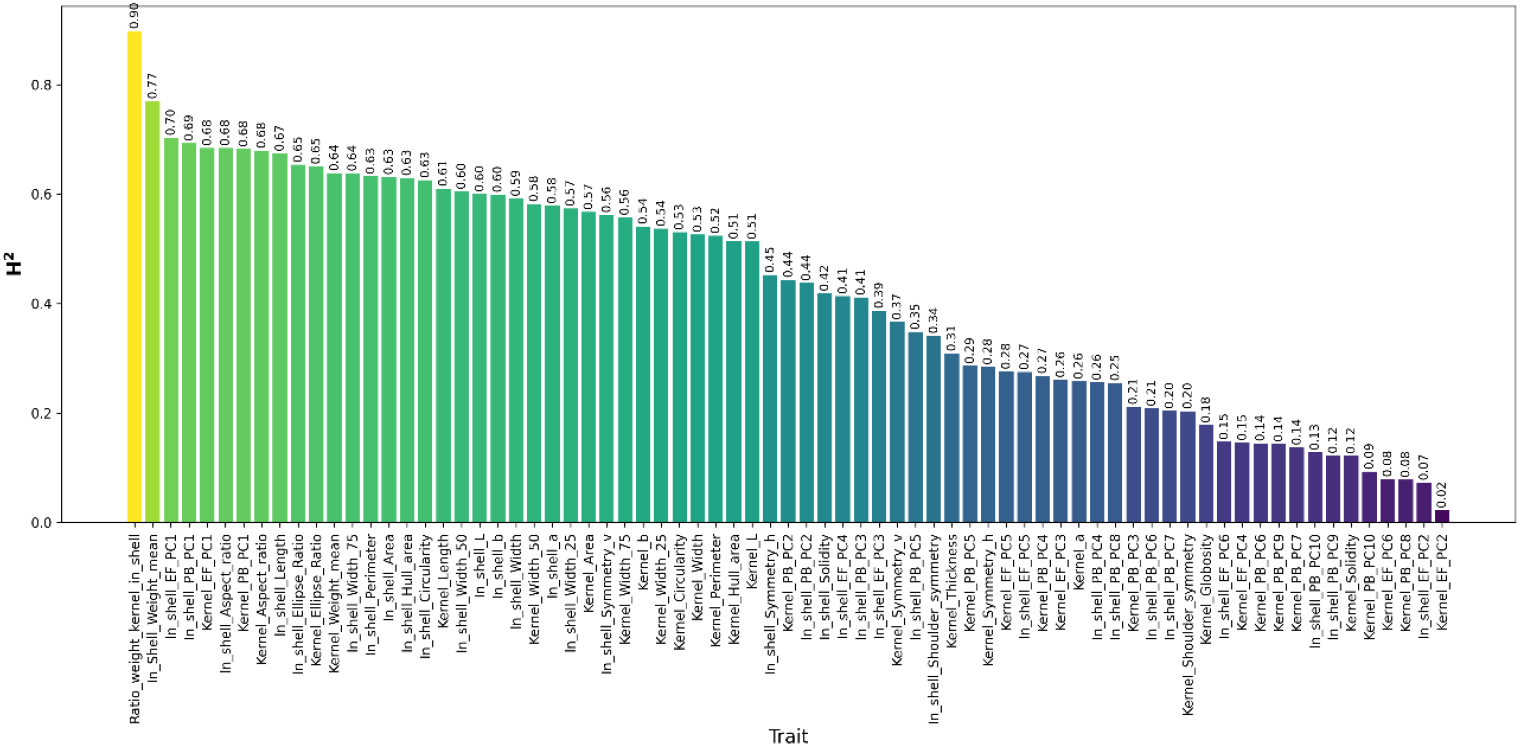
Broad-sense heritability of all traits studied in the almond populations.

**Figure 8.**
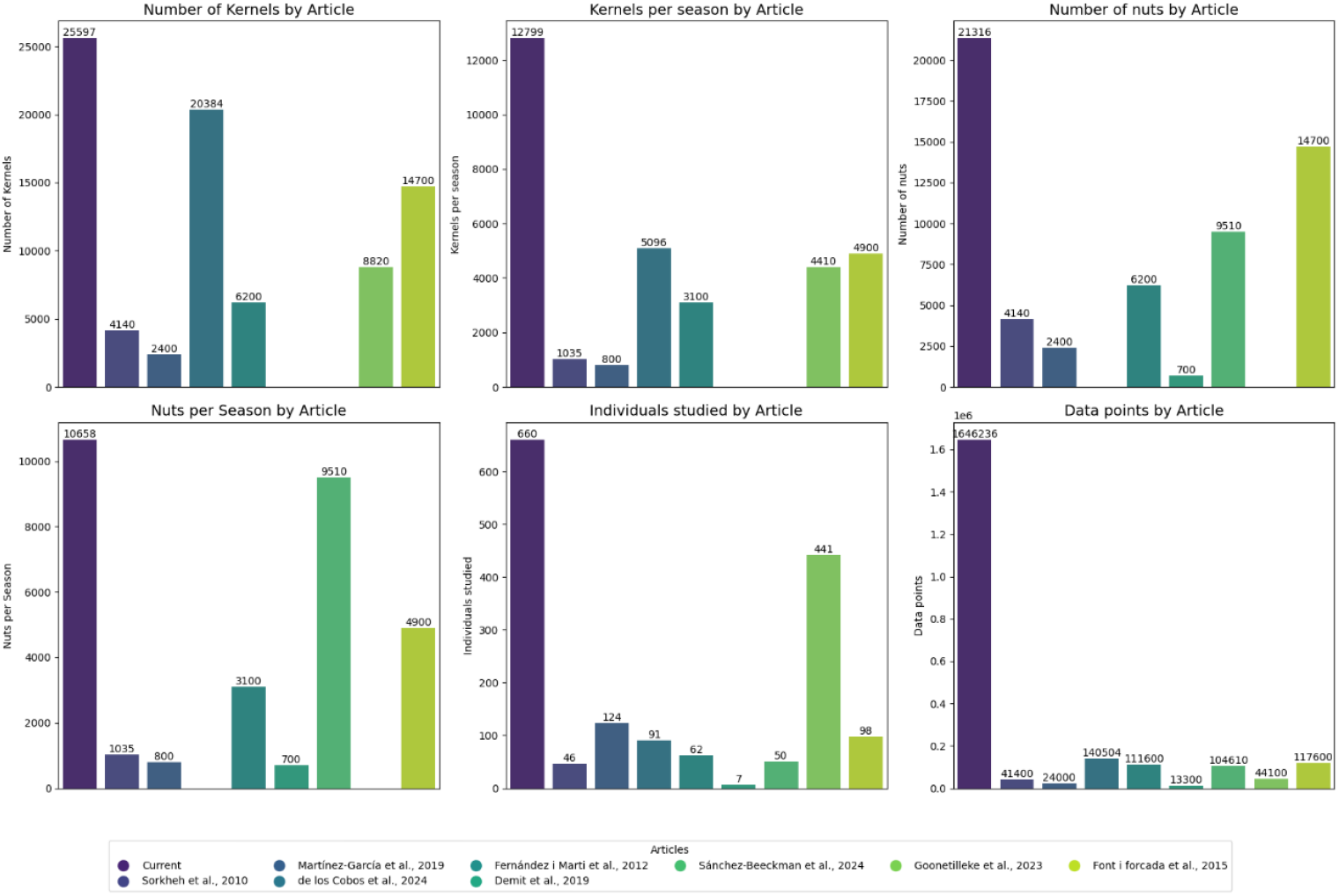
Bar plot for comparison among articles related to quantitative almond morphology traits (Demir et al., 2018; Fernández i Martí et al., 2013; Forcada et al., 2015; Goonetilleke et al., 2023; Martínez-García et al., 2019; Pérez de Los Cobos et al., 2024; Sánchez-Beeckman et al., 2024; Sorkheh et al., 2010).

### 3.5 Impact on almond breeding

Nine scientific articles related to almond morphology and breeding were collected to compare the performance of the high-throughput approach implemented in the current study (Supplementary Table 1 and Figure 8). In all the metrics analysed, the current study exceeded the phenotyping data collected in previous studies, with data points being 11 to 123 times higher than those reported in other works.

## 4. Discussion

The comprehensive workflow presented here introduces a new high-throughput phenotyping tool using almond as a case of study for fruits. This approach has enabled the analysis of 665 genotypes—the largest dataset for almond morphological quantitative traits reported to date, to the best of the authors’ knowledge. The development of this new phenotyping tool marks a major breakthrough in addressing the phenotyping bottleneck in almond. The developed models were adjusted to the target samples used but can be adapted to phenotype other kinds of fruits and other type of organs such as leaves or roots and more importantly as an open source, can be used by other plants breeding programs. In fact, the workflow was successfully applied to publicly available image datasets of apples and strawberries (Feldmann et al., 2020; Keller et al., 2024). Example results can be found in the workflow’s GitHub repository: https://github.com/jorgemasgomez/almondcv2.

Clearly, recent advancements in AI segmentation models, such as YOLO (Redmon et al., 2016) and SAM (Kirillov et al., 2023), enable breeding programs to develop fine-tuned models for specific applications, even without large datasets. Additionally, progress in labeling tools like CVAT (Sekachev et al., 2020), which integrate AI-assisted features, accelerates the tedious and time-consuming labeling process. The new workflow designed here uses Jupyter notebooks for ease of customization, eliminating the need for high-performance hardware (e.g., GPUs), allowing users to run the tool efficiently in cloud-based environments such as Google Colab. Moreover, the reconstruction approaches employed showed good performance in general terms. A lower performance was noticed only in the in-shell 2022 using SAHI which may be improved with SAHI parameter optimization.

The thickness estimation models demonstrated high accuracy, highlighting the usefulness of combining imaging phenotyping tools with traditional phenotyping processes, such as weighing (Utai et al., 2019). This approach enhances the phenotyping process by offering an alternative to capturing transversal pictures and makes phenotyping more accessible without relying on more complex instruments like 3D cameras or approaches as 3D reconstruction (Feldmann & Tabb, 2022; Sánchez-Beeckman et al., 2024). However, the approach employed takes advantage of the similar density among kernels, but it would not be effective for whole nuts due to the variability in shell hardness.

Morphometric analyses provided new quantitative traits for almond shape breeding. Both Elliptical Fourier Analysis and Pixel-based PCA showed a large variance explained by the PC1 in kernel and in-shell datasets. A high correlation between the different PCs1 and aspect ratio traits was identified and can also be observed graphically (Figure 3 and Supplementary Figure 7). Indeed, aspect ratio has been highly correlated with PCs1 in tomato and apple leaves (Chitwood et al., 2013; Migicovsky et al., 2018), in pear fruit (Wang et al., 2024), walnut (Demir et al., 2018), and in a shape discriminator almond nut (Demir et al., 2019). Although the interpretation of some morphometric traits could be abstract, patterns in different parts of the shape can be observed as in the tip (e.g. kernel PB-PC4), top (e.g. kernel PB-PC5) and side curvature (e.g. kernel EF-PC3). Moreover, some morphometric traits (apart from those related to aspect ratio) showed medium heritability values (>0.3) which could place them as breeding targets.

The number of kernels and nuts analysed in this comparative study is considerably higher than in previous research, and the difference becomes even more pronounced when comparing metrics per season. More importantly, this progress has been achieved while reducing both time and economic resources, as a single person was able to phenotype the entire dataset for one season in just two weeks. Currently, shell cracking remains the primary bottleneck due to the manual process, which is challenging to automate because of variability in shell hardness and size. This extensive dataset will facilitate future studies aimed at dissecting quantitative traits and implementing genomic selection approaches.

GPU: Graphics Processing Unit
YOLO: You Only Look Once
SAM: Segment Anything Model
ROI: Region of Interest

## Acknowledgements

This work was supported by grant PID2021-127421OB-I00 funded by MICIU/AEI/10.13039/501100011033, by grant CNS2022-135936 funded by MICIU/AEI /10.13039/501100011033 and European Union NextGenerationEU/PRTR and by “ERDF A way of making Europe” and formed part of the AGROALNEXT program and was supported by MCIN with funding from European Union NextGenerationEU (PRTR-C17.I1) and by the Fundación Séneca with funding from Comunidad Autónoma Región de Murcia (CARM). We would like to thank María del Mar Gómez Abajo, Francisco Gómez Lopez, Lucía Rodríguez Robles, Antonio Moreno Marín, Luis Miguel Serrano Sánchez, and Teresa Cremades Rosado for their assistance with the sampling, as well as Juan Antonio Tudela for lending us his camera and set. J.M.-G acknowledges the Spanish MICIU for his predoctoral grant (FPU20/00614).

