## Supplementary material for "Open RGB Imaging Workflow for Morphological and Morphometric Analysis of Fruits using AI: A Case Study on Almonds": Sup_Figures

[a]Fruit Breeding Group. Department of Plant Breeding, Centro de Edafología y Biología Aplicada del Segura- Spanish National Research Council (CEBAS-CSIC). Campus Universitario Espinardo, E-30100 Murcia, Spain

### Supplementary Figures

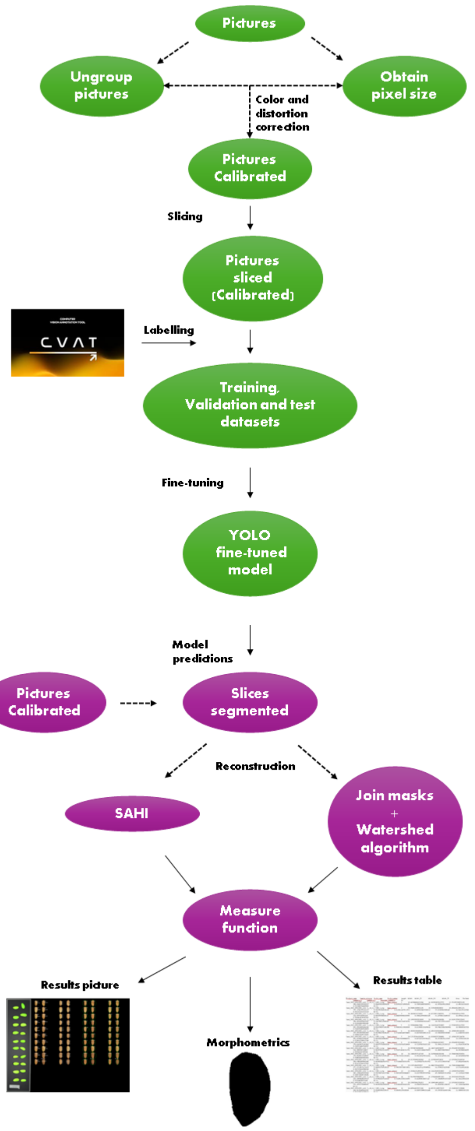

**Supplementary Figure 1.** Workflow description outlining the steps involved in developing the segmentation model (green) and deploying it (purple).

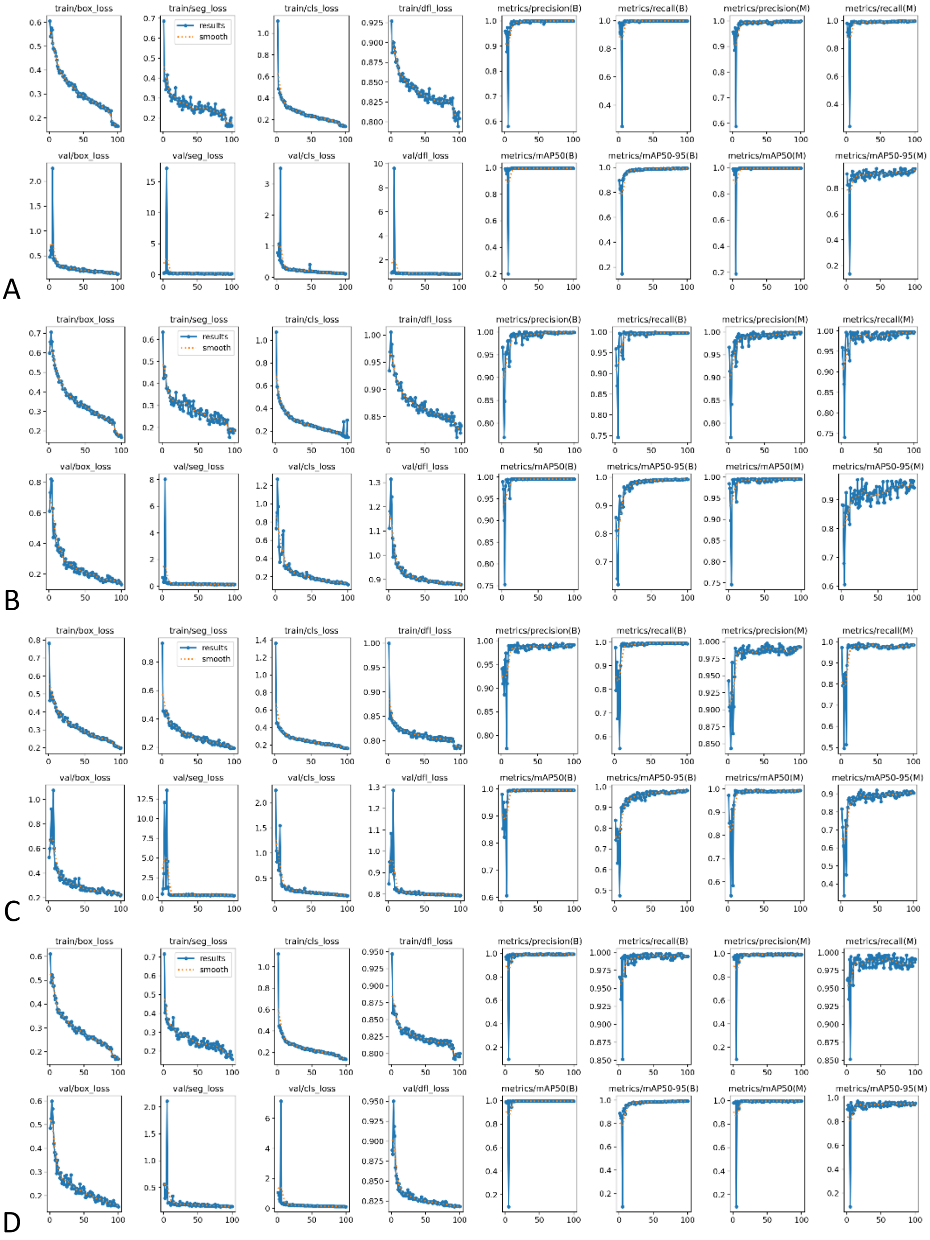


**Supplementary Figure 2**. YOLO metrics obtained during the fine-tuning process for the Kernel-2022 (A), In-shell-2022 (B), Kernel-2023 (C), and In-shell-2023 (D) datasets. A description of the metrics can be found in (Ultralytics, 2025).

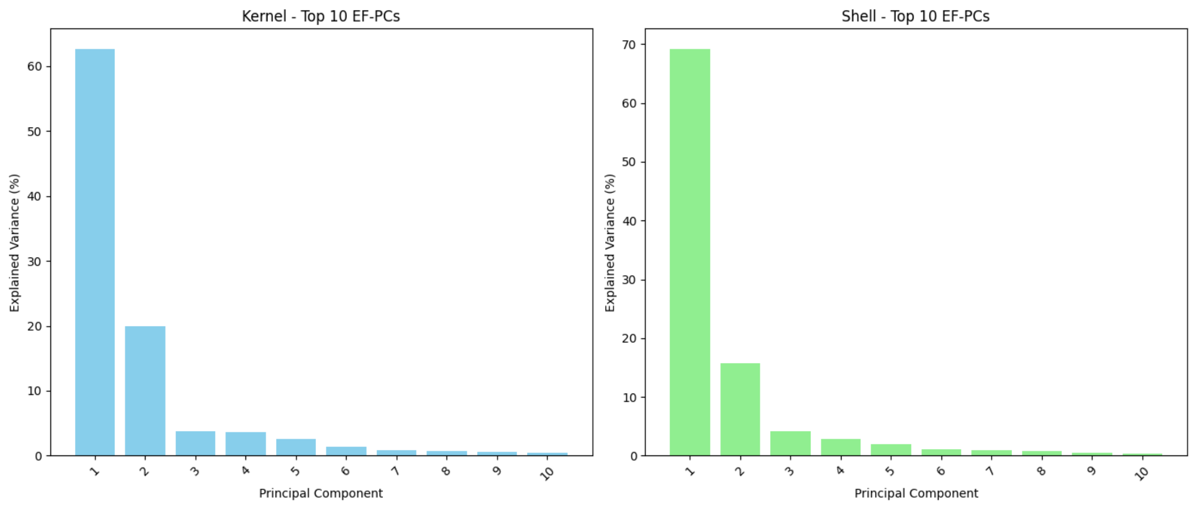


**Supplementary Figure 3.** Explained variance (%) per EF-PC in the kernel (left) and in-shell (right) datasets.

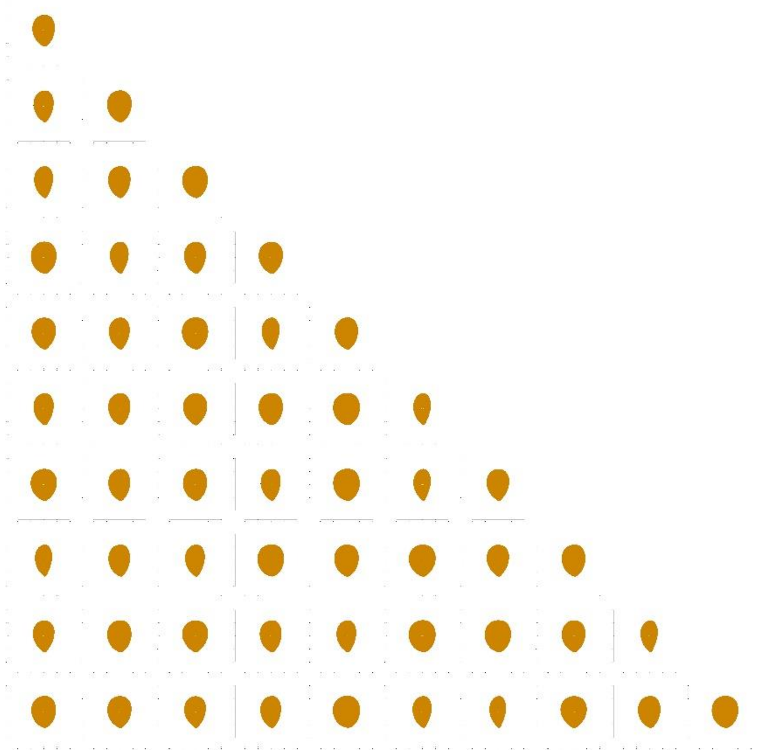

**Supplementary Figure 4.** K-means clustering using EFA-PCA results in in-shell dataset, showing the shapes corresponding to each centroid for scenarios ranging from k=1 to k = 10.

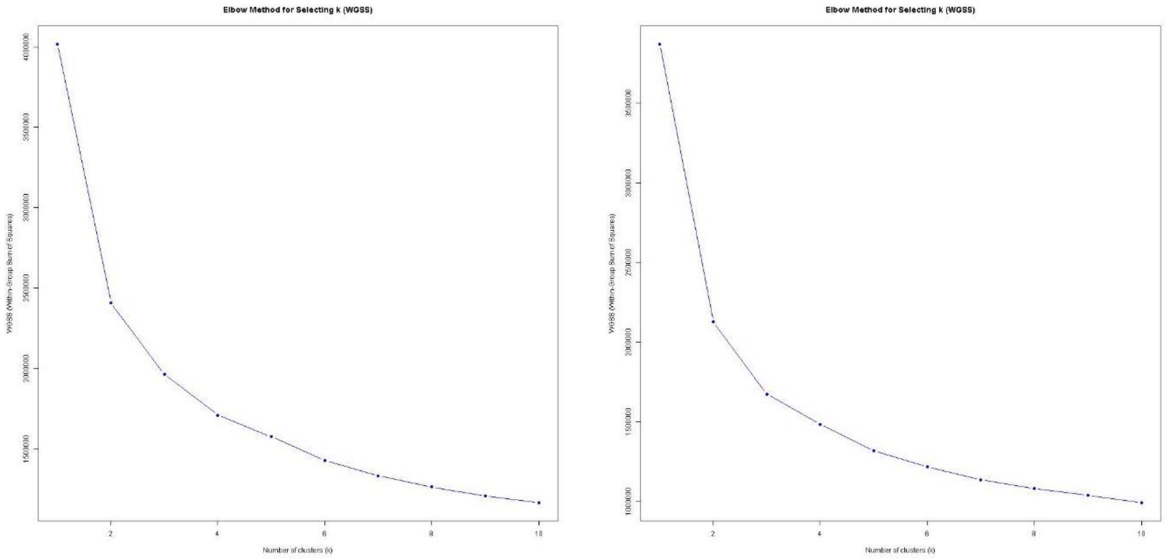


**Supplementary Figure 5.** Within group sum of squares decay for K-means clustering using EFA-PCA results in kernel (left) and in-shell (right) dataset.

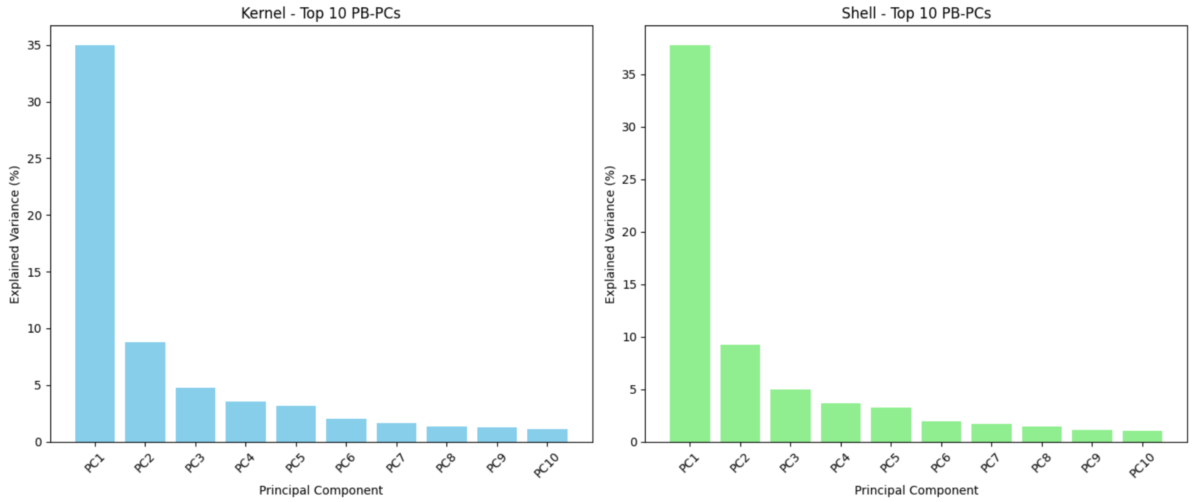


**Supplementary Figure 6.** Explained variance (%) per PB-PC in the kernel (left) and in-shell (right) datasets.

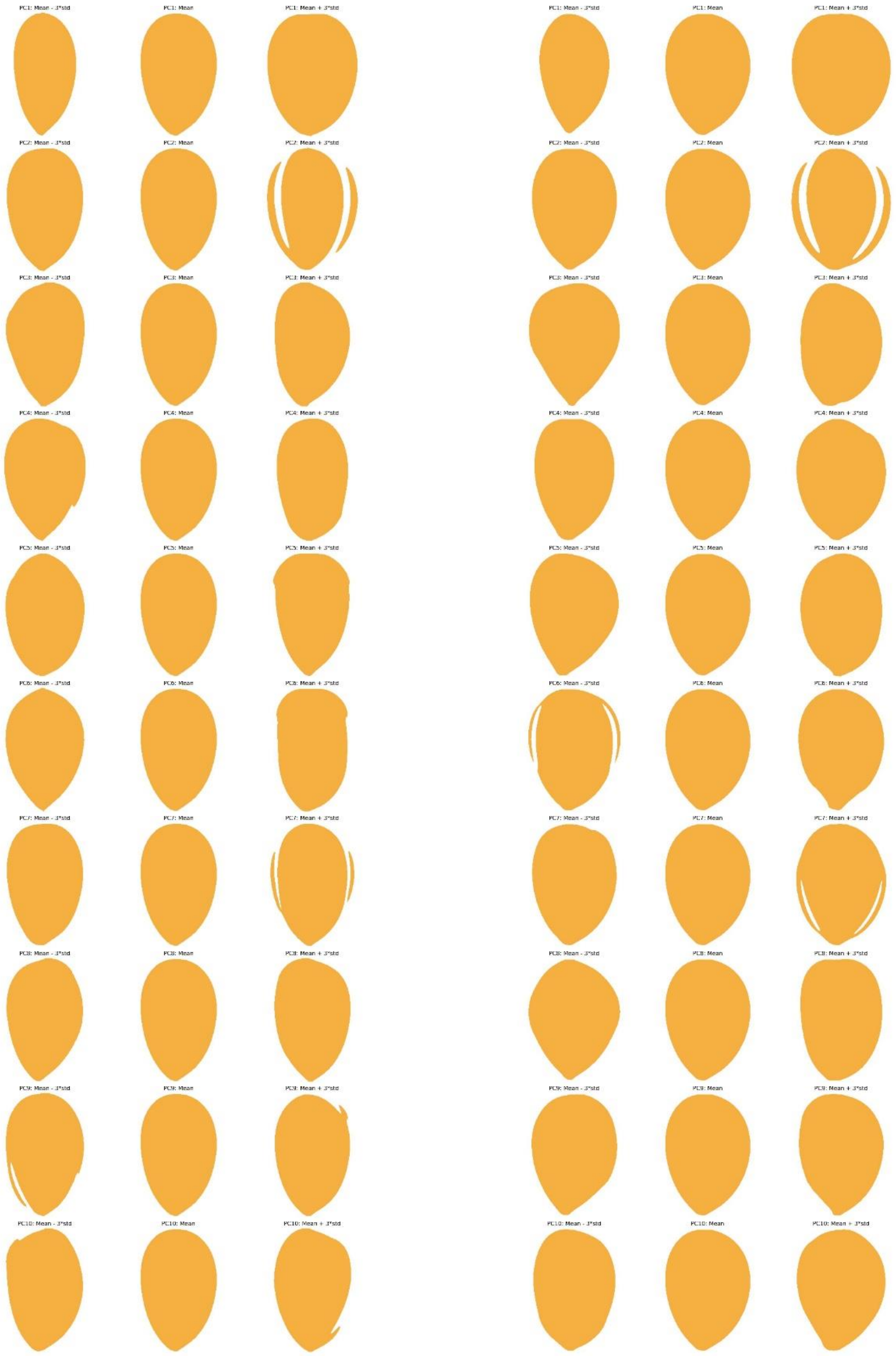


**Supplementary Figure 7.** Representation of the influence of the PB-PCs on shape in the kernel (left) and in-shell (right) datasets, from the mean shape to ±3× standard deviation.

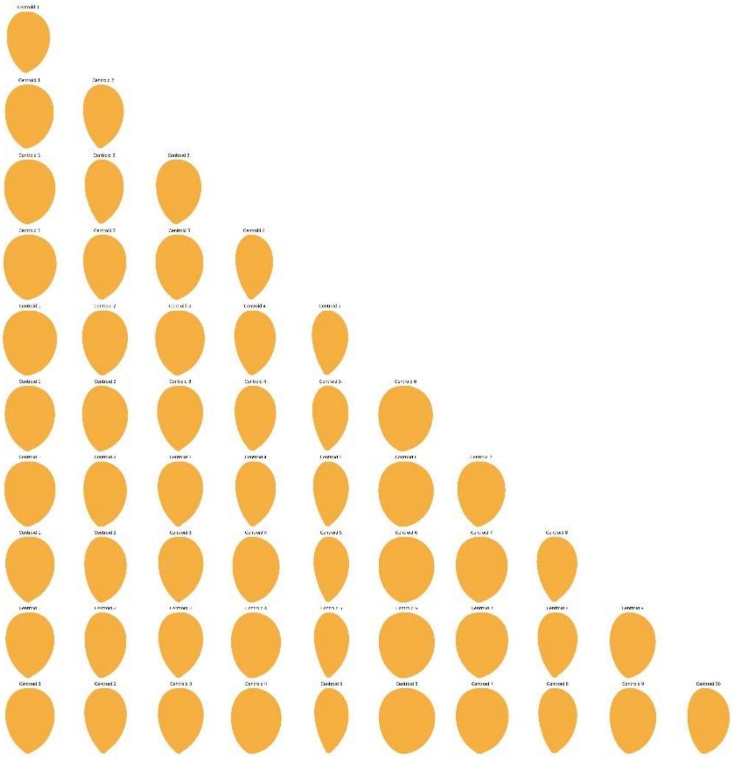

**Supplementary Figure 8.** K-means clustering using PB-PCA results in in-shell dataset, showing the shapes corresponding to each centroid for scenarios ranging from k=1 to k = 10.

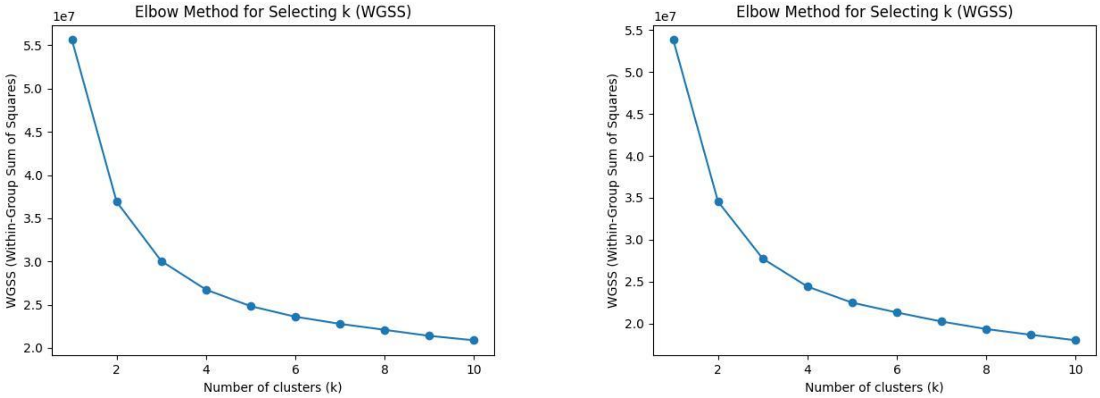


**Supplementary Figure 9.** Within group sum of squares decay for K-means clustering using PB-PCA results in kernel (left) and in-shell (right) dataset.
